# Tricalbin-mediated contact sites control ER curvature to maintain plasma membrane integrity

**DOI:** 10.1101/578427

**Authors:** Javier Collado, Maria Kalemanov, Antonio Martínez-Sánchez, Felix Campelo, Wolfgang Baumeister, Christopher J. Stefan, Ruben Fernández-Busnadiego

## Abstract

Membrane contact sites (MCS) between the endoplasmic reticulum (ER) and the plasma membrane (PM) play fundamental roles in all eukaryotic cells. ER-PM MCS are particularly abundant in *S. cerevisiae*, where approximately half of the PM surface is covered by cortical ER (cER). Several proteins, including Ist2, Scs2/22 and Tcb1/2/3 are implicated in cER formation, but the specific roles of these molecules are poorly understood. Here we use cryo-electron tomography to show that ER-PM tethers are key determinants of cER morphology. In particular, Tcb proteins form peaks of extreme curvature on the cER membrane facing the PM. Semi-quantitative modeling and functional assays suggest that Tcb-mediated cER peaks facilitate the transport of lipids from the cER to the PM, necessary to maintain PM integrity under stress conditions. ER peaks were also present at other MCS, implying that membrane curvature enforcement may be a widespread mechanism to expedite lipid transport at MCS.

## Introduction

Endoplasmic reticulum (ER)-plasma membrane (PM) membrane contact sites (MCS) are critical modulators of Ca^2+^ and lipid homeostasis in eukaryotic cells (Balla, 2018; Chang et al., 2017; Cockcroft and Raghu, 2018; Saheki and De Camilli, 2017a; Stefan, 2018). These structures, where the ER and the PM come into close apposition (10-30 nm), mediate store-operated Ca^2+^ entry (Carrasco and Meyer, 2011), insulin secretion by pancreatic beta cells (Lees et al., 2017) and excitation-contraction coupling in striated muscle (Bers, 2002). Consequently, dysregulation of ER-PM MCS is linked to multiple human diseases (Lacruz and Feske, 2015; Landstrom et al., 2014; Rios et al., 2015).

ER-PM MCS are particularly abundant in the yeast *Saccharomyces cerevisiae*. In these cells, a large portion of the ER is found at the cell cortex forming MCS with the PM. The extent of these contacts is such that nearly half of the PM surface area is covered by cortical ER (cER) (Manford et al., 2012; Pichler et al., 2001; Quon et al., 2018; Toulmay and Prinz, 2012; West et al., 2011). The loss of six proteins (Ist2, Scs2/22 and Tcb1/2/3; “Δtether” cells) dramatically reduces the extent of ER-PM association, indicating that these proteins are important ER-PM tethers in *S. cerevisiae* (Manford et al., 2012). Additional proteins, including Ice2 and the yeast StARkin orthologs, are also implicated in cER-PM function (Gatta et al., 2015; Quon et al., 2018). Loss of cER triggers PM lipid imbalance (Manford et al., 2012; Quon et al., 2018), highlighting the physiological importance of these membrane junctions.

Ist2 is a member of the anoctamin/TMEM16 protein family (Whitlock and Hartzell, 2017). Ist2 resides on the ER membrane and consists of eight transmembrane domains plus a long C-terminal cytoplasmic tail that binds PM lipids (Figure S 1), thereby tethering the ER and the PM (Fischer et al., 2009; Juschke et al., 2005; Maass et al., 2009; Manford et al., 2012). Deletion of Ist2 results in reduced cER levels, whereas Ist2 overexpression leads to increased ER-PM MCS (Manford et al., 2012; Wolf et al., 2012).

Scs2/22 are orthologues of the mammalian VAMP-associated proteins (VAPs), a family of ER-resident proteins widely implicated in MCS formation (Murphy and Levine, 2016; Stefan et al., 2011). Both Scs2 and Scs22 are C-terminally anchored to the ER by a transmembrane segment, and contain an N-terminal major sperm protein domain (MSP) (Figure S 1). Scs2/22 function as ER-PM tethers thanks to the binding of their MSP domain to PM proteins containing FFAT or FFAT-like motifs (Manford et al., 2012; Murphy and Levine, 2016). A strong reduction in cER levels is observed in Scs2/22 knockout (KO) cells (Loewen et al., 2007; Manford et al., 2012).

The tricalbin proteins (Tcb1/2/3) are orthologues of the mammalian extended-synaptotagmins (E-Syts) and the plant synaptotagmins (SYTs) (Perez-Sancho et al., 2016; Saheki and De Camilli, 2017b). Tcbs are likely anchored to the ER membrane by a hairpin sequence (Giordano et al., 2013; Saheki and De Camilli, 2017b) (Figure S 1) similar to those found in ER morphogenetic proteins such as reticulons (Hu et al., 2011). Tcbs harbor a synaptotagmin-like, mitochondrial and lipid binding protein (SMP) domain that can bind and transport lipids (Lee and Hong, 2006; Saheki et al., 2016; Schauder et al., 2014; Toulmay and Prinz, 2012; Yu et al., 2016). SMP domains have been found in multiple MCS-resident proteins, and likely play a key role in the inter-membrane exchange of lipids at these sites (Reinisch and De Camilli, 2016). C-terminal to the SMP domain, Tcbs contain a variable number of C2 domains (four in Tcb1/2 and five in Tcb3), some of which can bind membrane phospholipids in a manner either dependent or independent of Ca^2+^ (Creutz et al., 2004; Rizo and Sudhof, 1998; Schulz and Creutz, 2004). Both the SMP and C2 domains are required for Tcb targeting to the cER (Manford et al., 2012; Toulmay and Prinz, 2012), and tethering likely takes place via PM binding by C2 domains (Giordano et al., 2013).

Although Ist2, Scs2/22 and Tcb1/2/3 are involved in the appropriate formation of cER, the exact functions of these proteins at ER-PM MCS are poorly understood. First, whereas Ist2 and Scs2/22 are important ER-PM tethers, their relative contributions to cER generation remain unclear. The functions of Tcbs are even more mysterious: Tcbs are bona fide ER-PM tethers, because cER levels in mutants expressing Tcbs but lacking Ist2 and Scs2/22 are significantly higher than in Δtether cells (Manford et al., 2012). However, loss of Tcbs on their own does not result in a substantial reduction in the amount of cER (Manford et al., 2012; Toulmay and Prinz, 2012), suggesting that the main role of Tcbs is not the mechanical anchoring of the ER to the PM. More broadly, the physiological functions of the mammalian E-Syts remain similarly unclear (Sclip et al., 2016; Tremblay and Moss, 2016), although their capacity to shuttle lipids at ER-PM MCS has been demonstrated (Bian et al., 2018; Saheki et al., 2016; Yu et al., 2016).

Here we aimed to dissect the functional roles of Ist2, Scs2/22 and Tcb1/2/3 at ER-PM MCS. Toward this end, we used cryo-electron tomography (cryo-ET) to study the fine structure of the cER within mutant cells lacking specific tethers. Thanks to the advent of cryo-focused ion beam (cryo-FIB) technology and direct electron detectors, cryo-ET allows high resolution 3D imaging of a virtually unperturbed cell interior at molecular resolution (Beck and Baumeister, 2016; Rigort et al., 2012; Wagner et al., 2017). Given the minute dimensions of MCS, these structures are particularly sensitive to alterations introduced by classical EM procedures such as chemical fixation, dehydration and heavy metal staining. Therefore, cryo-ET is especially suited for the high-resolution study of native MCS architecture (Collado and Fernandez-Busnadiego, 2017; Fernandez-Busnadiego et al., 2015). Our results show that, besides simply anchoring the ER to the PM, each family of tethers uniquely contributes to shaping the cER. In particular, Scs2/22 are associated with cER sheets, whereas Tcbs favor cER tubules. Notably, Tcbs are necessary for the generation of peaks of extreme curvature at the cER membrane that contribute to maintaining PM integrity, possibly by facilitating the transport of cER lipids to the PM.

## Results

### MCS Architecture in *S. cerevisiae*

To study MCS architecture *in situ* by cryo-ET, *S. cerevisiae* cells were vitrified on EM grids and thinned down to 100-200 nm-thick lamellae using cryo-FIB. Lamellae were loaded into a cryo-TEM (Figure S 2), and tomograms were acquired at suitable cellular locations. Cryo-tomograms of various MCS (Figure 1A, B) revealed abundant proteinaceous densities of diverse morphologies bridging the gap between the membranes (Figure 1C, D; Figure S 3A). Interestingly, distance measurements showed a characteristic inter-membrane spacing for different MCS. For example, while average nucleus-vacuole distance was 21 ± 7 nm (mean ± STD, N = 5 nucleus-vacuole MCS; Figure 1E), ER-mitochondria junctions were significantly narrower (16 ± 7 nm; mean ± STD, N = 6 ER-mitochondria MCS; p < 0.05 by unpaired t-test; Figure 1E). These data suggest that the optimal distance range for inter-organelle communication is different at different MCS.

**Figure 1:**
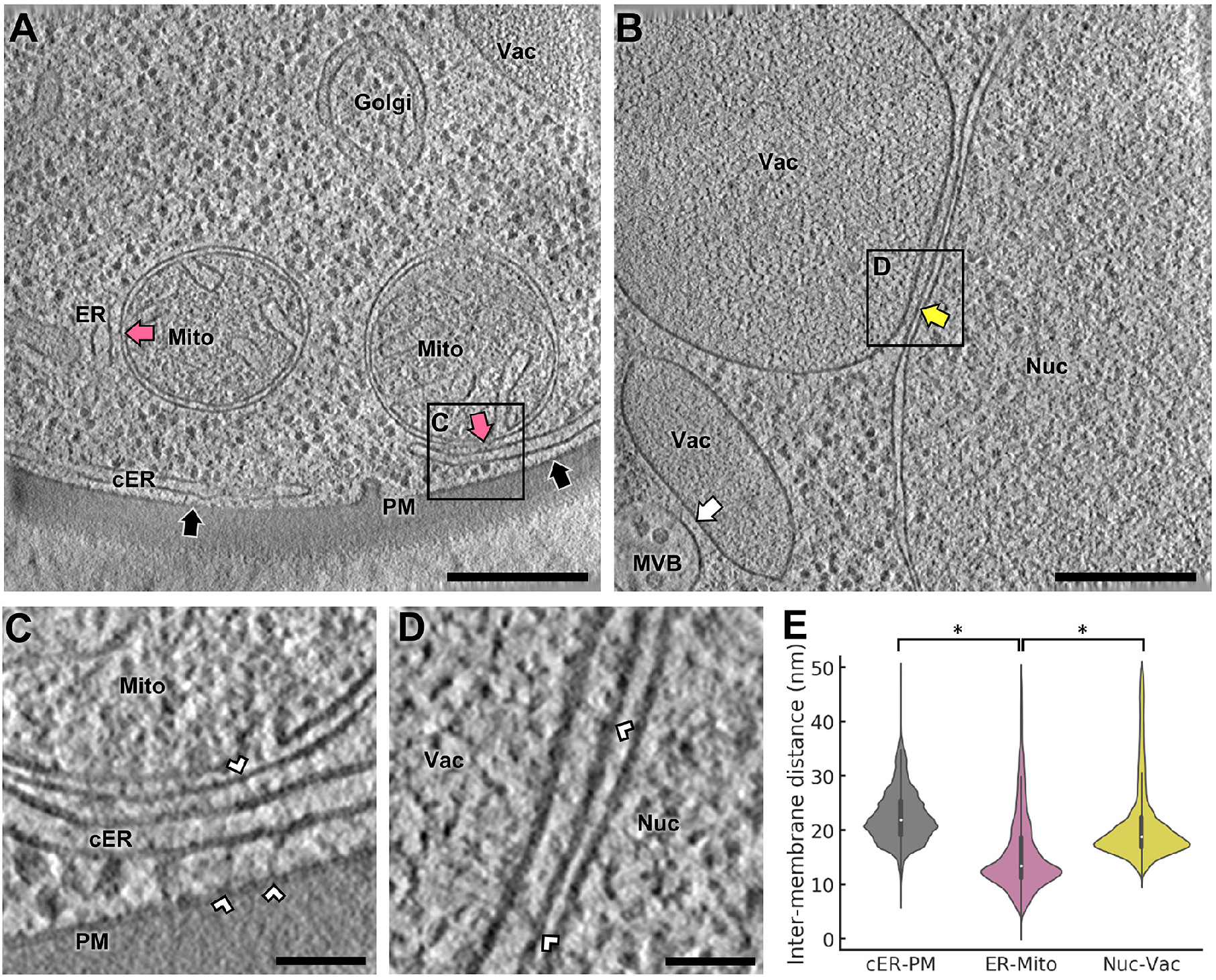
Cryo-ET Imaging of MCS in WT *S. cerevisiae*. **(A)** 1.4 nm-thick tomographic slice showing cER-PM MCS (black arrows) and ER-mitochondria MCS (purple arrows). The boxed area is magnified in (C). ER: endoplasmic reticulum; cER: cortical ER; Golgi: Golgi apparatus; Mito: mitochondrion; PM: plasma membrane; Vac: vacuole. **(B)** 1.4 nm-thick tomographic slice showing a nucleus-vacuole junction (yellow arrow) and a multivesicular body-vacuole MCS (white arrow). The boxed area is magnified in (D). MVB: multivesicular body; Nuc: nucleus. **(C)** Magnification of the area boxed in (A). White arrowheads: inter-membrane tethers. **(D)** Magnification of the area boxed in (B). **(E)** Violin plots showing the distribution of inter-membrane distances of cER-PM, ER-mitochondrion and nucleus-vacuole MCS. * indicates p < 0.05 by unpaired t-test. N = 5 (cER-PM), 6 (ER-mitochondria) and 5 (nucleus-vacuole) MCS in WT cells. Scale bars: 300 nm (A, B), 50 nm (C, D). See also Figure S 3.

To gain further insights into the molecular determinants of MCS structure and function we focused on ER-PM MCS, perhaps the most prominent MCS in *S. cerevisiae* (Manford et al., 2012; Pichler et al., 2001; Quon et al., 2018; West et al., 2011). Cryo-ET analysis showed that the cER of WT cells consisted of both membrane sheets and tubules (Figure 2A), with an average thickness of 25 ± 6 nm (mean ± STD, N = 5 cER-PM MCS; Figure 2F). For 95 % of the MCS area, ER-PM distance ranged from 15 to 33 nm, with an average of 22 ± 4 nm (mean ± STD; Figure 1E, Figure 2E). Thus, in WT cells the cER had a variable morphology and a relatively broad distribution of distances to the PM.

**Figure 2:**
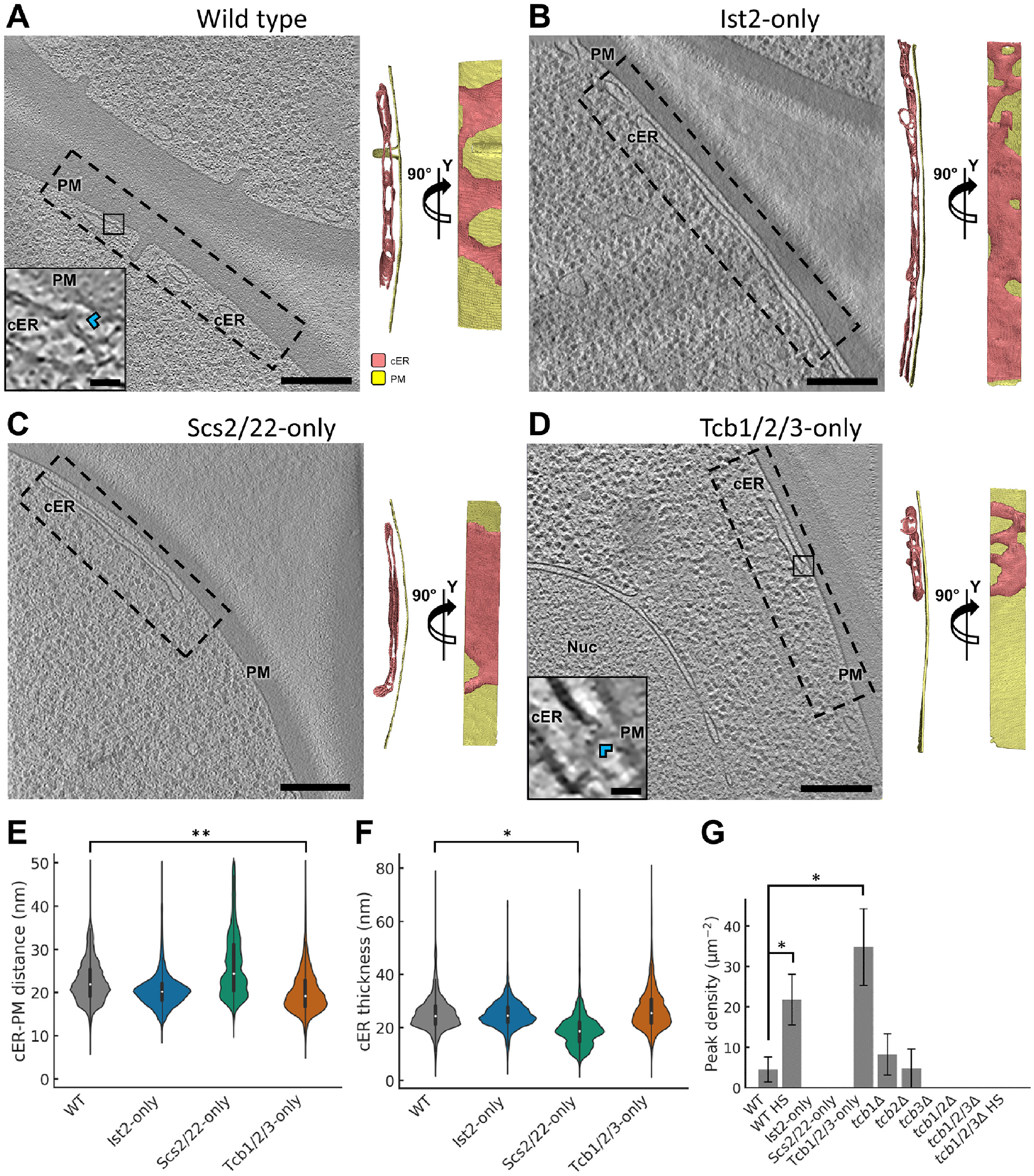
cER Morphology in ER-PM MCS Tether Mutants. Panels (A-D) show 1.4 nm-thick tomographic slices of cER in the indicated strains (left) and 3D renderings in two perpendicular orientations (right). cER: cortical ER (pink); Nuc: nucleus; PM: plasma membrane (gold). **(A)** WT cell, **(B)** Ist2-only cell, **(C)** Scs2/22-only cell, **(D)** Tcb1/2/3-only cell. Insets in (A) and (D) show cER peaks (blue arrowheads). Scale bars: 300 nm (main panels), 25 nm (insets). Panels (E-G) show quantifications of cER-PM distance **(E)**, cER thickness **(F)** and cER peak density per μm^2^ of cER membrane area **(G)**. HS: heat shock (42 °C for 15 min). * and ** indicate respectively p < 0.05 and p < 0.01 by unpaired t-test. N = 5 (WT), 7 (WT HS), 4 (Ist2-only), 4 (Scs2/22-only), 6 (Tcb1/2/3-only), 4 *(tcblΔ)*, 4 *(tcb2Δ)*, 4 *(tcb3Δ)*, 4 *(tcb1/2Δ)*, 4 *(tcb1/2/3Δ)*, and 4 *(tcb1/2/3Δ* HS) cER-PM MCS. See also Figure S 1, Figure S 2 and Figure S 3.

### ER-PM Tethers Control cER Morphology

Because the simultaneous deletion of Ist2, Scs2/22 and Tcb1/2/3 largely abolishes ER-PM MCS (Manford et al., 2012), we sought to understand the individual contribution of each of these molecules to ER-PM tethering. To that end, we performed cryo-ET imaging of ER-PM MCS in mutant cells expressing only one family of tethers. These data confirmed previous observations (Loewen et al., 2007; Manford et al., 2012; Toulmay and Prinz, 2012; Wolf et al., 2012) that total levels of cER were still substantial in cells expressing only Ist2 *(scs2/22Δ tcb1/2/3Δ;* “Ist2-only” cells; Figure S 2B) or the VAP orthologues Scs2 and Scs22 *(ist2Δ tcb1/2/3Δ;* “Scs2/22-only” cells; Figure S 2C). However, cER levels in cells expressing only Tcb1/2/3 *(ist2Δ scs2/22Δ;* “Tcb1/2/3-only” cells; Figure S 2D) were markedly lower than in WT (Figure S 2A), although higher than in Δtether cells (Figure S 2E), in agreement with previous results (Manford et al., 2012).

Next, we investigated the fine morphology of the cER in these mutants. In Ist2-only cells, the cER was a mixture of membrane sheets and tubules similar to WT cells (Figure 2A, B). Average ER-PM distance (21 ± 4 nm, mean ± STD, N = 4 cER-PM MCS; Figure 2E) and cER thickness (25 ± 6 nm, mean ± STD; Figure 2F) were also comparable to WT, suggesting that Ist2 is an important contributor to the morphology of the cER in WT cells.

Interestingly, cER morphology in Scs2/22-only cells was dramatically different from WT. In these mutants cER tubules were rarely observed, as the cER consisted almost exclusively of extended sheets (Figure 2C). Average ER-PM distance in these cells spread across a wider range of values (26 ± 8 nm, mean ± STD, N = 4 cER-PM MCS; Figure 2E). On the other hand, the ER sheets observed in Scs2/22-only cells were significantly narrower than WT (19 ± 6 nm, mean ± STD; p < 0.05 by unpaired t-test; Figure 2F). These data show that while Scs2/22 are not very effective in controlling ER-PM distance, they are important determinants of cER width.

In contrast to Scs2/22-only cells, the cER was formed mainly by membrane tubules in Tcb1/2/3-only cells (Figure 2D). Interestingly, in these cells we also observed abundant peaks of very high curvature on the membrane of the cER facing the PM (Figure 2D, inset; Figure S 3B). These peaks had a diameter of ~20 nm and protruded 5-10 nm from the cER membrane (Figure S 3C). At cER peaks the cER came into very close proximity of the PM, reaching distances of ~7 nm (Figure S 3C). Overall, the average cER-PM distance was significantly shorter in Tcb1/2/3-only (20 ± 5 nm, mean ± STD; N = 6 cER-PM MCS; p < 0.01 by unpaired t-test; Figure 2E) than in WT cells. Whereas cER peaks were not found in Ist2-only (Figure 2B, G) or Scs2/22-only (Figure 2C, G) cells, they were also present in WT cells, albeit at lower frequency (Figure 2A, inset; Figure 2G; Figure S 3B). cER peaks in WT cells were morphologically indistinguishable from those found in Tcb1/2/3-only cells (Figure S 3C). Therefore, the formation of the high curvature cER peaks in WT cells may be controlled by the Tcb proteins. Because the peaks were only found on the part of the cER membrane opposed to the PM, these structures may be involved in inter-membrane crosstalk.

Altogether, these data show that ER-PM tethers play a key role in controlling cER morphology, especially in terms of membrane curvature.

### Quantitative Analysis of cER Membrane Curvature

Membrane curvature plays a major role in a wide variety of cellular processes (Kozlov et al., 2014) and is a fundamental determinant of ER morphology (Hu et al., 2011). Therefore, we further analyzed the cER membrane curvature alterations observed in the different ER-PM tether mutants. To that end, we implemented an algorithm allowing quantitative determination of membrane curvature in cryo-ET data (Kalemanov et al., 2019). A global analysis was consistent with the visual impression that the average cER curvature observed in Scs2/22-only cells was lower than in WT (p < 0.01 by unpaired t-test; Figure 3A, C, E), indicating the higher prevalence of cER sheets. Conversely, the curvature of the cER membrane in Tcb1/2/3-only cells was significantly higher than WT (p < 0.01 by unpaired t-test; Figure 3A, D, E), reflecting the more tubular cER morphology in these cells. Local mapping of the curvature in cER membrane renderings highlighted the presence of peaks of extreme curvature (curvature radius ≤ 10 nm; Figure 3A, inset) in WT cells. These structures were enriched in Tcb1/2/3-only cells compared to WT (p < 0.05 by unpaired t-test; Figure 2G; Figure 3D, inset), and absent in *tcb1/2/3Δ* mutants (N = 4 cER-PM MCS; Figure 2G; Figure 4E) and cells expressing only Scs2/22 or Ist2 (Figure 2G; Figure 3B, C). Therefore, this analysis reinforced the notion that Tcb1/2/3 are necessary for the generation of high curvature peaks on the side of the cER membrane directly facing the PM.

**Figure 3:**
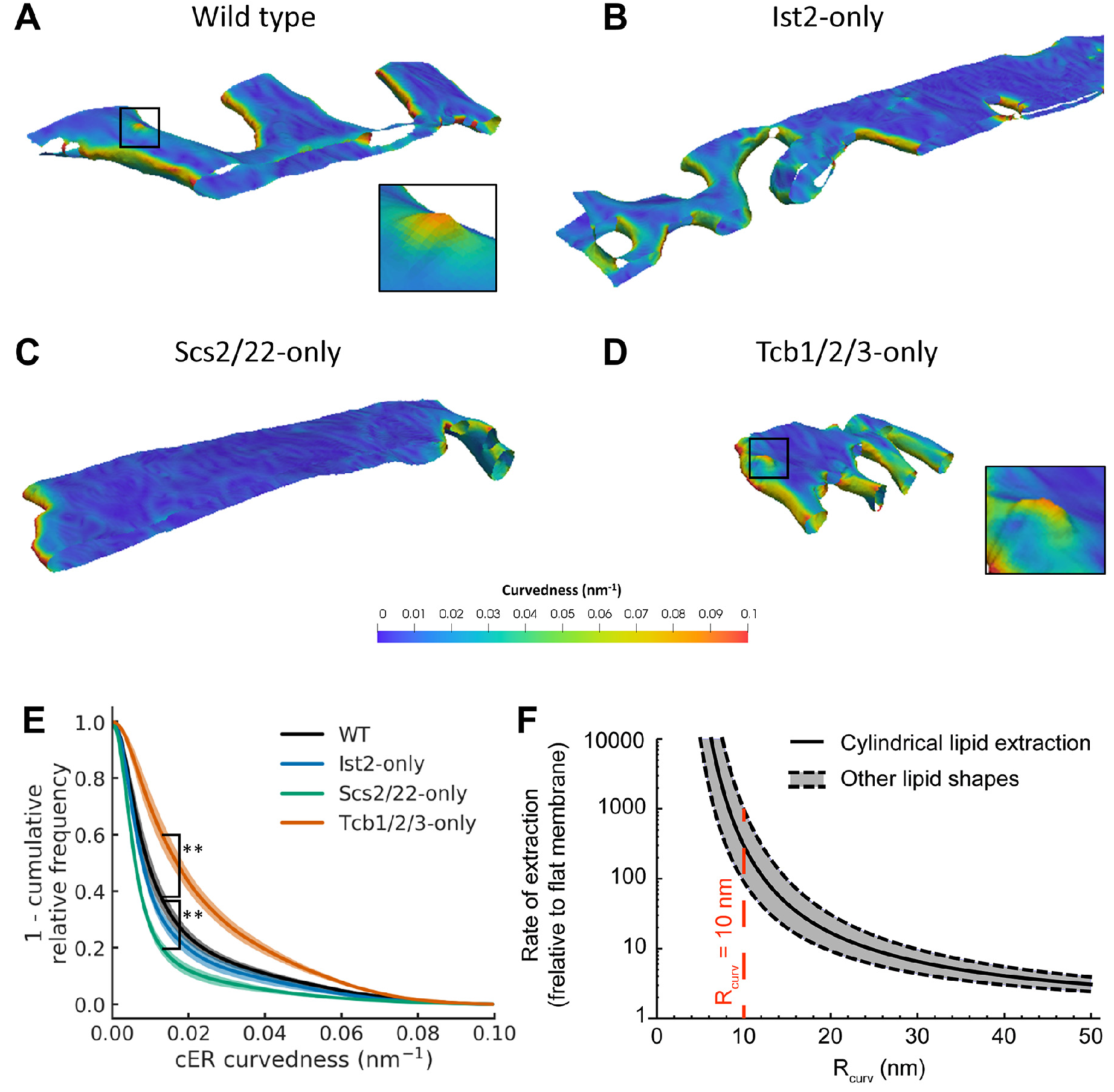
Quantification of cER Curvature. Panels (A-D) show 3D visualizations of cER curvedness in the indicated strains. Insets in (A) and (D) show cER peaks. **(A)** WT cell, **(B)** Ist2-only cell, **(C)** Scs2/22-only cell, **(D)** Tcb1/2/3-only cell. **(E)** Quantification of cER curvedness. ** indicates p < 0.01 by unpaired t-test. N = 5 (WT), 7 (WT HS), 4 (Ist2-only), 4 (Scs2/22-only), and 6 (Tcb1/2/3-only) cER-PM MCS. **(F)** Enhancement of the rate of lipid extraction by membrane curvature according to a theoretical model. The plot shows the rate of extraction computed for a standard cylindrical lipid (black curve) as well as for lipids of other shapes, such as conical or inverted conical lipids (gray-shaded area between the dashed, black curves). The value of the radius of curvature of the experimentally observed cER peaks is denoted by the dashed red line. 1/*R_curv_* is equivalent to the curvedness for k_1_=k_2_. See also Figure S 4.

To address the molecular basis of this phenomenon, we next investigated which Tcb proteins were required for cER peak formation by conducting cryo-ET of cells lacking specific Tcbs in the presence of all other ER-PM tethers. In *tcb1Δ* (N = 4 cER-PM MCS; Figure 4A; Figure S 3B) and *tcb2Δ* cells (N = 4 cER-PM MCS; Figure 4B; Figure S 3B), cER peaks were observed at a similar frequency than in WT cells (Figure 2G). cER peaks in these strains were also comparable to WT in terms of diameter, height and distance to the PM (Figure S 3C). In contrast, cER peaks were extremely rare in *tcb3Δ* (N = 4 cER-PM MCS; Figure 2G; Figure 4C) and *tcb1/2Δ* cells (N = 4 cER-PM MCS; Figure 2G; Figure 4D). Therefore, the expression of Tcb3 and either Tcb1 or Tcb2 seems necessary for the efficient formation of cER peaks, consistent with reports that Tcbs (and E-Syts) can form heterodimers (Creutz et al., 2004; Giordano et al., 2013; Idevall-Hagren et al., 2015).

**Figure 4:**
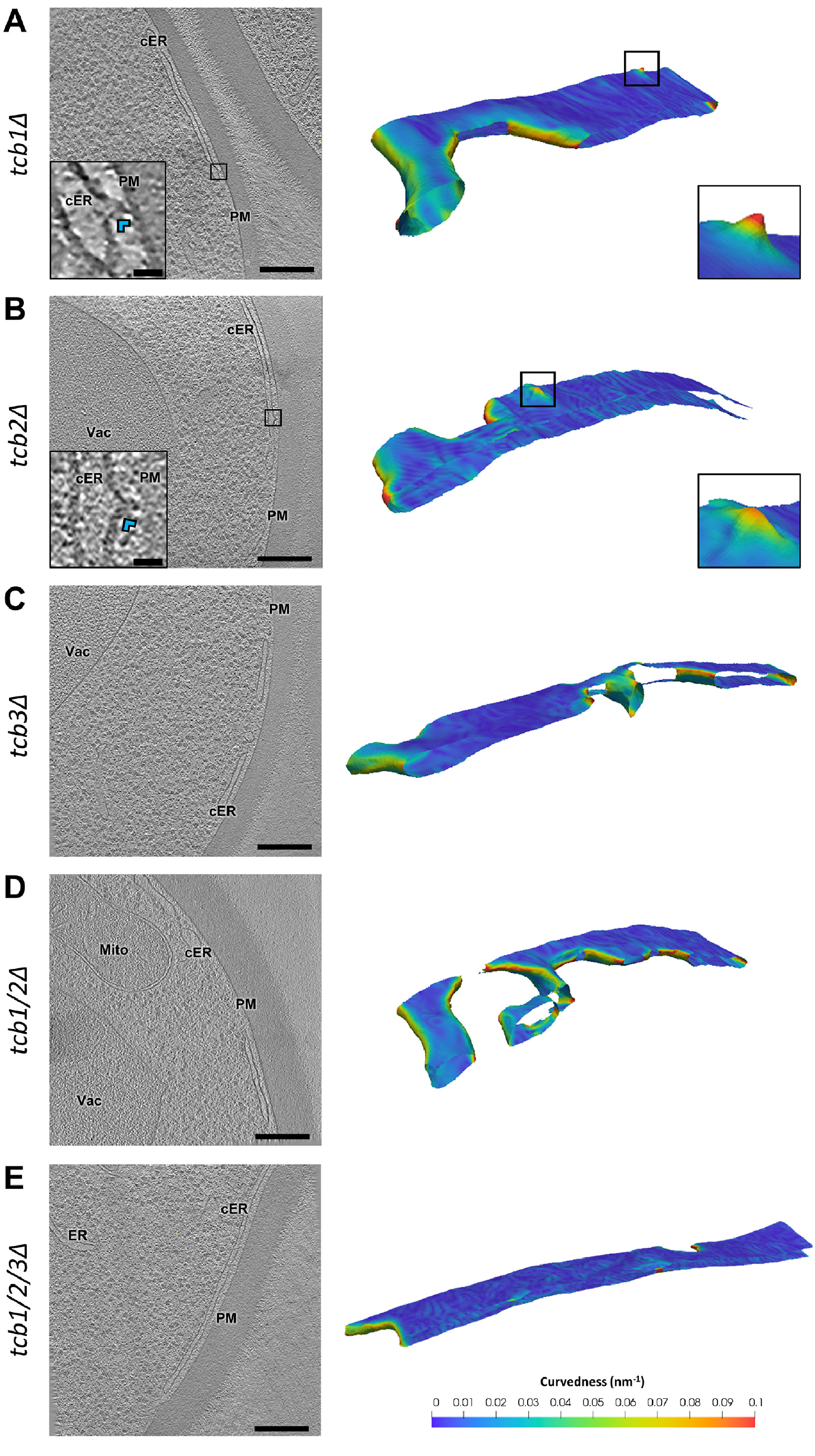
cER Peaks in Tcb Mutants. Panels (A-E) show 1.4 nm-thick tomographic slices of cER in the indicated strains (left) and 3D renderings of cER curvature (right). **(A)** *tcblΔ*, **(B)** *tcb2Δ*, **(C)** *tcb3Δ*, **(D)** *tcb1/2Δ*, **(E)** *tcb1/2/3Δ* cell. cER: cortical ER; Mito: mitochondrion; PM: plasma membrane; Vac: vacuole. Insets in (A) and (B) show cER peaks (blue arrowheads). Scale bars for tomographic slices: 300 nm (main panels), 25 nm (insets). See also Figure S 3.

### cER Peaks May Facilitate Lipid Transfer

Next, we investigated the biological function of Tcb-mediated cER peaks. Tcbs contain modules that can sense or induce membrane curvature (hairpin anchor, multiple C2 domains) and transport lipids (SMP domain) (Creutz et al., 2004; Lee and Hong, 2006; Manford et al., 2012; Toulmay and Prinz, 2012). Tcbs may combine both their curvature generation and lipid transport properties by controlling the formation of cER peaks, which could facilitate cER-to-PM lipid transport by i) reducing the physical distance between cER and PM, and/or ii) disturbing the cER lipid bilayer to facilitate lipid extraction, and at the same time impose cER-to-PM directionality to the transfer process.

To address this possibility, we used a semi-quantitative model (Campelo and Kozlov, 2014) to calculate how the induction of cER membrane curvature may facilitate the lipid transfer process. We assume that this task is performed by a lipid transport module such as the SMP domain of Tcbs. The total free energy required for lipid extraction by a lipid transport protein (LTP) can be expressed as the sum of two components. The first one is independent from membrane geometry, incorporating electrostatic interactions and membrane-independent interactions between the lipid and the LTP. The second component is determined by the elastic stresses imposed on the membrane by its geometry prior to LTP binding, and by how these stresses change as a result of the lipid rearrangements caused by a partial membrane insertion of the LTP. In turn, membrane geometry can be determined by its lipid composition and/or by external factors such as curvature-generating proteins (Campelo et al., 2008). We focused on this last scenario, as it is unlikely that physiological lipid compositions result in the extreme membrane curvatures of cER peaks (Campelo et al., 2008; Sorre et al., 2009). With these premises, our calculations showed that the energy barrier for lipid extraction is reduced by ~6 kBT when the radius of curvature of the membrane is 10 nm (Figure S 4), as observed in Tcb-induced cER peaks. This is of similar magnitude to the facilitation of sterol extraction from a flat membrane by an LTP in comparison to its spontaneous desorption, estimated to be ~2-3 k_B_T (Dittman and Menon, 2017), and would result in a 500-1000 fold acceleration of the transfer reaction (Figure 3F). Therefore, our model predicts that cER peaks greatly facilitate lipid extraction by lipid transport modules.

### cER Peaks Maintain PM Integrity

The synthesis of most PM lipids – including phosphatidylinositol, phosphatidylserine, and sterols– is enhanced at the cER (Pichler et al., 2001). Therefore, if cER peaks are important for the transport of newly synthesized lipids from the cER to the PM, cells lacking cER peaks may suffer from PM lipid imbalance. In line with this hypothesis, *tcb1/2/3Δ* cells show PM integrity defects upon heat stress (Omnus et al., 2016), a situation in which substantial delivery of lipids from the ER to the PM may be necessary to repair heat-induced alterations. Because Tcbs are required for the formation of cER peaks that may facilitate lipid transfer, it is possible that cER peaks are important for ER-PM lipid transport under heat stress conditions.

To test this hypothesis, we performed PM integrity assays in the different Tcb mutants. Cells were subjected to 42 °C for 15 min, and PM integrity was monitored by measuring the entry of extracellular propidium iodide into cells using flow cytometry (Figure 5A). Because this dye is membrane impermeable, it only enters cells with compromised PM integrity (Zhao et al., 2013). Remarkably, these experiments revealed PM integrity defects only for conditions in which cER peaks were not observed *(tcb3Δ, tcb1/2/3Δ* cells; p < 0.01 by unpaired t-test; Figure 5A), whereas PM integrity in Tcb mutants showing cER peaks *(tcb1Δ, tcb2Δ)* was comparable to WT (Figure 5A). Thus, there was a strong correlation between the absence of Tcb-induced cER peaks and PM integrity defects.

**Figure 5:**
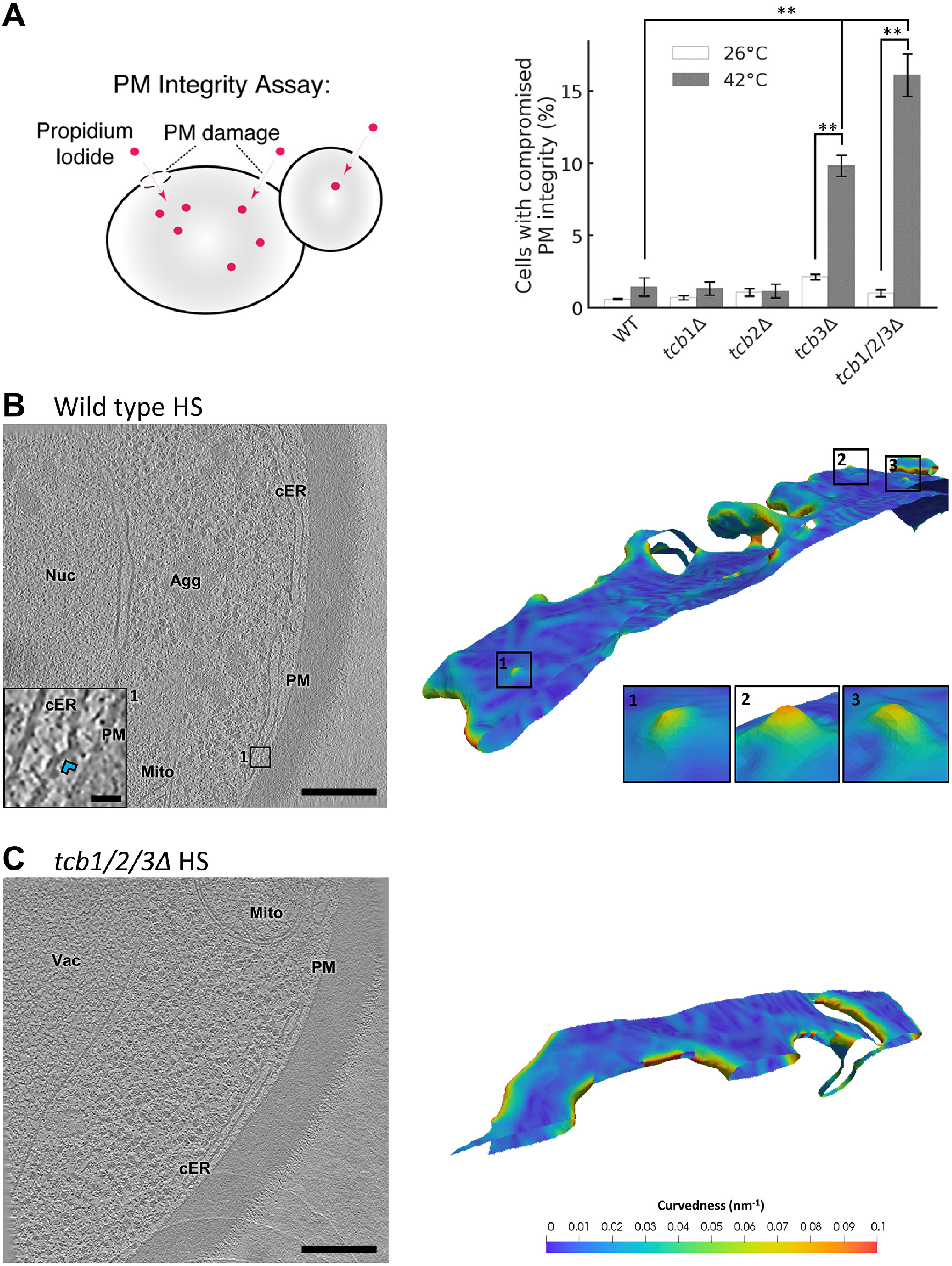
PM Integrity and cER Morphology under Heat Stress. **(A)** Schematic of the propidium iodide assay to assess PM integrity (left) and PM integrity measurements of Tcb deletion mutants upon 15 min incubation at 42 °C (right). The entry of propidium iodide in cells with compromised PM integrity was measured by flow cytometrxy. Three independent biological repeats were performed for all conditions. Panels (B-C) show 1.4 nm-thick tomographic slices of cER in the indicated strains (left) and 3D renderings of cER curvature (right). Agg: aggregate; cER: cortical ER; Mito: mitochondrion; Nuc: nucleus; PM: plasma membrane; Vac: vacuole. **(B)** WT cell under heat stress (HS). Insets show cER peaks (blue arrowhead in the tomographic slice inset). **(C)** *tcb1/2/3Δ* cell under heat stress. Scale bars: 300 nm (main panels), 25 nm (inset). See also Figure S 5.

However, the density of cER peaks was relatively low in WT cells under non-stress conditions (Figure 2A, G; Figure 3A). Could these rare structures play an important role in maintaining PM integrity under heat shock conditions? To address this question, we performed cryo-ET on heat-shocked cells. As expected, cells showed abundant amorphous aggregates in various cellular locations (Miller et al., 2015; Wagner et al., 2017)(Figure 5B). Also, WT cells showed a strong increase in the number of cER peaks (N = 6 cER-PM MCS; p < 0.05 by unpaired t-test; Figure 2G; Figure 5B), whereas these structures were absent in heat-shocked *tcb1/2/3Δ* cells (N = 4 cER-PM MCS; Figure 2G; Figure 5C). Therefore, the Tcb-dependent formation of cER peaks is induced by conditions that challenge PM integrity, such as heat shock.

Altogether, these data indicate that i) Tcbs are necessary for the formation of cER peaks, ii) cER peaks may facilitate ER-to-PM lipid transfer, and iii) cER peaks are important to maintain PM integrity under heat stress, a condition in which the PM may require substantial lipid flux to restore its lipid homeostasis. Interestingly, similar high curvature peaks were observed at other ER-mediated MCS such as ER-mitochondria MCS (Figure S 3D), suggesting that the induction of membrane curvature is a general mechanism to facilitate inter-membrane lipid exchange at various MCS.

## Discussion

### ER-PM Tethers Shape the cER

MCS are now known to exist between essentially all cellular membranes, and a great number of MCS-resident proteins and tethers have been identified. However, the functions of many of these molecules remain poorly understood (Bohnert and Schuldiner, 2018; Shai et al., 2018; Valm et al., 2017; Wu et al., 2018). For example, it is unclear why six proteins (Ist2, Scs2/22 and Tcb1/2/3) are necessary to anchor the ER to the PM in *S. cerevisiae* (Manford et al., 2012), as well as the possible functions of these proteins beyond the mechanical stapling of the membranes. Here, we employed state-of-the-art *in situ* imaging by cryo-ET to reveal that ER-PM tethers are critical determinants of cER morphology and MCS function.

Our data confirms that Ist2 is an important ER-PM tether (Lavieu et al., 2010; Manford et al., 2012; Wolf et al., 2012). The distribution of ER-PM distances was particularly narrow in Ist2-only cells, indicating that Ist2 is very effective in maintaining an ER-PM separation of about 21 nm. This is surprising, because Ist2 bridges the ER and the PM by a 340 amino acid-long linker that is predicted to be unstructured, and could thus extend up to 120 nm. Interestingly, Kralt et al. proposed that shorter linkers that should still be able to span a ~20 nm distance if unstructured (240, 140 or 58 amino acids long) effectively result in much shorter ER-PM distances, as suggested by their localization to progressively smaller puncta (Kralt et al., 2015). How a presumably disordered 340 amino acid stretch can precisely regulate ER-PM inter-membrane distance to 21 nm requires further investigation.

ER-PM distances were much more broadly distributed in Scs2/22-only cells, possibly due to the promiscuous interactions of Scs2/22 with different FFAT/FFAT-like motif proteins at the PM (Murphy and Levine, 2016). Strikingly, the cER in these cells consisted almost exclusively of extended, narrow sheets. How such sheets are formed remains to be established, as direct interactions of Scs2/22 in trans across the ER lumen appear unlikely, given their short luminal sequences. In contrast to Scs2/22-only cells, the cER was mainly formed by tubules in Tcb1/2/3-only cells. This phenomenon may rely on the hairpin sequence that anchors Tcbs to the ER membrane, which could sense and/or generate membrane curvature as in reticulons and other ER morphogenetic proteins (Hu et al., 2011).

Alternatively, it is also possible that the cER morphologies observed in the different ER-PM tether mutants arise from dysregulation of additional ER-PM MCS factors (Quon et al., 2018), other ER morphogenetic proteins or lipids. However, WT cER also consists of a mixture of sheets and tubules, which could plausibly arise as a combination of the morphologies of the individual ER-PM tether mutants. Therefore, it is possible that the different families of tethers are enriched in partially segregated cER subdomains, consistent with their punctate localization observed by light microscopy (Manford et al., 2012; Toulmay and Prinz, 2012; Wolf et al., 2012). Similarly, in plant and mammalian cells different ER-PM tethers are known to co-exist at the same MCS but form separate subdomains (Giordano et al., 2013; Siao et al., 2016). Therefore, native ER-PM MCS may be established as a juxtaposition of molecular territories enriched in different tethers.

### Tcb-Mediated cER Peaks May Facilitate ER-to-PM Lipid Transfer

Besides being generally tubular, the cER in Tcb1/2/3-only cells was enriched in membrane peaks of extreme (<10 nm radius) membrane curvature, which were also present in WT cells at a lower frequency. Therefore, it is likely that Tcbs generate the high curvature peaks observed in the cER of WT cells. We speculate that this action is carried out by the binding of Tcb C2 domains to the cER membrane, by analogy with other multi-C2 domain proteins. For example, the C2 domains of mammalian SYT1 bind the PM in a Ca^2+^-dependent manner, generating curvature in the PM to reduce the energy barrier for synaptic vesicle fusion (Martens et al., 2007). The exact mechanisms by which Tcb C2 domains could generate such peaks, possibly involving Tcb oligomerization (Creutz et al., 2004; Giordano et al., 2013; Zanetti et al., 2016), require further investigation.

Tcb-induced cER peaks always faced the PM, suggesting roles in inter-membrane exchange. Unlike synaptic vesicles and the PM, the ER and the PM are not thought to fuse (Saheki and De Camilli, 2017a). However, most MCS harbor an extensive non-vesicular exchange of lipids (Cockcroft and Raghu, 2018; Lees et al., 2017; Saheki et al., 2016), especially important at ER-PM MCS because most PM lipids are synthesized in the ER. Importantly, our semi-quantitative modeling indicates that cER peaks can dramatically enhance the transfer of lipids from the cER to the PM by facilitating the shallow insertion of lipid transport modules into the lipid bilayer, in agreement with experimental studies (Moser von Filseck et al., 2015). cER peaks also shorten cER-PM distance, and can impose cER-to-PM directionality to the lipid transfer process. Altogether, cER-to-PM lipid transfer may be greatly enhanced by Tcb-mediated cER peaks.

Supporting this hypothesis, the E-Syts, mammalian orthologues of Tcbs, are also implicated in ER-PM lipid transfer (Bian et al., 2018; Saheki et al., 2016; Yu et al., 2016). Yeast Tcbs, plant SYTs and mammalian E-Syts contain an SMP domain that mediates lipid binding and transport (Bian et al., 2018; Saheki et al., 2016; Schauder et al., 2014; Yu et al., 2016). It is attractive to speculate that these proteins combine two actions to catalyze cER-to-PM lipid transport: i) C2 domain-mediated generation of extreme curvature in the cER to substantially lower the energy barrier for lipid extraction, and ii) SMP domain-mediated lipid binding and subsequent transport to the PM.

Bona fide membrane curvature generators such as reticulons have also been implicated in ER-PM MCS formation (Caldieri et al., 2017), and the activity of ER-PM MCS-resident lipid synthesizing enzymes may be regulated by membrane curvature (Bozelli et al., 2018). Furthermore, there is increasing evidence for important roles of curvature sensing/generating proteins at other MCS (Ackema et al., 2016; de Saint-Jean et al., 2011; Ho and Stroupe, 2016; Moser von Filseck et al., 2015; Voss et al., 2012), consistent with our observations of high curvature peaks at e.g. ER-mitochondria MCS. Thus, membrane curvature may be an important regulator of MCS function (Henne et al., 2015).

### cER Peaks Are Important for PM Integrity under Stress

The physiological roles of the Tcb protein family remain enigmatic. On one hand, these molecules are highly conserved and therefore likely to play important functions. However, no major alterations were discovered in yeast cells lacking all Tcbs (Manford et al., 2012; Toulmay and Prinz, 2012), nor in mammalian cells or mice lacking all three E-Syts (Saheki et al., 2016; Sclip et al., 2016; Tremblay and Moss, 2016). E-Syt triple knockout cells did display an accumulation of diacylglycerol at the PM upon phospholipase C activation (Saheki et al., 2016), suggesting that the main function of E-Syts/Tcbs is to respond to stimuli that perturb lipid homeostasis (Stefan, 2018).

Our data suggest that one of such stimuli is heat stress. Although Tcb1/2/3 deletion does not substantially reduce the levels of cER, upon heat stress *tcb1/2/3Δ* cells suffer from PM integrity defects similar to Δtether cells (Omnus et al., 2016). These defects are not rescued by an artificial linker, indicating a specific requirement for Tcbs. In line with this idea, our functional assays of different Tcb mutants showed a strong correlation between the absence of cER peaks and PM integrity defects upon heat stress. Although the exact mechanisms by which heat stress compromises PM integrity remain to be established, heat alters PM protein and lipid homeostasis, as well as the physico-chemical properties of the bilayer (Fan and Evans, 2015; Verghese et al., 2012; Zhao et al., 2013). PM repair likely involves the addition of new lipids (Vaughan et al., 2014), and ER-PM MCS regulate phospholipid biogenesis (Pichler et al., 2001; Tavassoli et al., 2013). Thus, under conditions of PM damage, Tcb-mediated cER peaks could maintain PM integrity by ensuring sufficient flow of lipids newly synthesized in the cER towards the PM.

Consistently, we observed a substantial increase in the number of cER peaks in WT cells under heat stress. Since membrane binding of some Tcb C2 domains is regulated by Ca^2+^ (Schulz and Creutz, 2004), the formation of new cER peaks upon heat stress may be driven by the influx of extracellular Ca^2+^ through a damaged PM (Andrews and Corrotte, 2018; Jimenez and Perez, 2017). Ca^2+^ influx may trigger the binding of Tcb C2 domains to the cER membrane, inducing cER membrane curvature in a way similar to mammalian SYTs (Martens et al., 2007). At the same time, the cER is highly dynamic, and explores most of the cellular PM over a few minutes, possibly monitoring PM status (Omnus et al., 2016). Thus, Ca^2+^ signals at sites of PM damage may trigger localized formation of Tcb-mediated cER peaks to locally enhance PM repair (Figure S 5). This mechanism may act in parallel or synergistically with other pathways to maintain PM homeostasis (Andrews and Corrotte, 2018; Jimenez and Perez, 2017; Omnus et al., 2016; Zhao et al., 2013).

These notions agree with findings on the plant SYTs, which also act as ER-PM tethers and have been extensively characterized as important factors for maintaining PM integrity under different stresses (Kawamura and Uemura, 2003; Lee et al., 2019; Perez-Sancho et al., 2015; Schapire et al., 2008; Yamazaki et al., 2008). Other mammalian multi-C2 domain proteins like SYT7 and dysferlin are directly implicated in plasma membrane repair (Andrews and Corrotte, 2018; Jimenez and Perez, 2017). Like yeast Tcbs, membrane binding by some C2 domains of plant SYTs and mammalian E-Syts is regulated by Ca^2+^ (Giordano et al., 2013; Idevall-Hagren et al., 2015; Perez-Sancho et al., 2015). Therefore, we propose that a crucial function of yeast Tcbs, plant SYTs and possibly mammalian E-Syts is to respond to influx of extracellular Ca^2+^ through a damaged PM by forming cER peaks, which drive a cER-to-PM lipid transfer necessary for PM repair. Further work should identify the lipid species involved in this inter-membrane exchange.

## Abbreviations

cER: cortical endoplasmic reticulum
cryo-ET: cryo-electron tomography
ER: endoplasmic reticulum
E-Syt: extended synaptotagmin
KO: knockout
LTP: lipid transport protein
MCS: membrane contact site
MSP: major sperm protein
PM: plasma membrane
SMP: synaptotagmin-like, mitochondrial and lipid binding protein
SYT: synaptotagmin
Tcb: tricalbin
VAP: VAMP-associated proteins

## Acknowledgments

We thank Günter Pfeifer, Jürgen Plitzko and Miroslava Schaffer for electron microscopy support, Scott Emr for strains, as well as Markus Hohle, Vladan Lucic, Eri Sakata and Florian Wilfling for helpful discussions. We also thank Patrick C. Hoffmann and Wanda Kukulski for sharing unpublished results. J.C. and M.K. are supported by the Graduate School of Quantitative Biosciences Munich. J.C., M.K., W.B. and R.F.-B. have received funding from the European Commission (FP7 GA ERC-2012-SyG_318987–ToPAG). F.C. acknowledges financial support from the Spanish Ministry of Economy and Competitiveness (“Severo Ochoa” program for Centres of Excellence in R&D (SEV-2015-0522), FIS2015-63550-R, FIS2017-89560-R, and BFU2015-73288-JIN, AEI/FEDER/UE), Fundació Privada Cellex and from the Generalitat de Catalunya through the CERCA program. C.S. is supported by MRC funding to the MRC LMCB University Unit at UCL, award code MC_UU_00012/6.

## Author Contributions

J.C. performed electron microscopy experiments and contributed to computational data analysis. M.K. and A.M.S. developed software procedures for data analysis. F.C. and M.F.G.-P. performed theoretical modeling. W.B., C.J.S. and R.F.-B. designed research. C.J.S. constructed strains and performed plasma membrane integrity assays. R.F.-B. supervised electron microscopy experiments and data analysis. R.F.-B. wrote the manuscript. All authors contributed to the manuscript.

## STAR Methods

### Yeast Strains and Cell Culture

The yeast strains used in this study are listed in Table S1.

Yeast colonies grown on YPD plates were inoculated in liquid YPD and incubated at 30 °C until reaching 0.6 OD_600_.

### Cell Vitrification

Cryo-EM grids (R2/1, Cu 200 mesh grid, Quantifoil Micro Tools) were glow discharged using a plasma cleaner (PDC-3XG, Harrick) for 30 s and mounted on a Vitrobot Mark IV (FEI).

A 3.5 μl drop of yeast culture was deposited on the carbon side of the grid before being blotted from the back using filter paper (Whattman 597) at force setting 9 for 10 s. The grids were immediately plunged into a liquid ethane/propane mixture at liquid nitrogen temperature and stored in sealed boxes submerged in liquid nitrogen until usage.

### Cryo-Focused Ion Beam Milling

Vitrified grids were mounted into Autogrid carriers (FEI), held in place by a copper ring. They were subsequently inserted in a dual-beam Quanta 3D cryo-FIB / scanning electron microscope (SEM) (FEI) using a transfer shuttle and a cryo-transfer system (PP3000T, Quorum). Inside the microscope the sample was kept at −180°C using a cryo-stage throughout the milling process.

To protect the sample from unwanted damage by the ion beam, a layer of organic platinum was deposited on top of the grid using a gas injection system from a 13.5 mm distance for 9 s. Small groups of cells located near the center of the grid square were targeted for milling. Milling was done at a 20° tilt. Several sequential steps were taken, starting with the Ga^2+^ ion beam at 30 kV and 500 pA beam current for rough milling, down to 30 kV and 30 pA for fine milling.

The final lamellae were around 14 μm wide and 150-250 nm thick. SEM imaging at 5 kV and 13 pA was used to monitor the milling process. The final thickness was reached when the lamella lacked contrast at 3 kV and 10 pA.

### Cryo-Electron Tomography

The lamellae were imaged at liquid nitrogen temperature in a Titan Krios cryo-electron microscope (FEI) equipped with a 300 kV field emission gun, a post column energy filter (Gatan) and a K2 Summit direct electron detector (Gatan).

Low magnification (3600 X, 40 Å pixel size) images of the lamellae were taken to identify regions of interest. Tilt series were recorded using SerialEM software (Mastronarde, 2005) at higher magnification (42,000 X, 3.42 Å pixel size), typically from −46° to +64° with increments of 2°. The camera was operated in dose fractionation mode, producing frames every 0.2 s. 1/cos scheme was used to increase the exposure time at higher tilt angles, resulting in exposure times of 1-2 s per projection image. Tilt series for *tcb1/2/3Δ*, WT heat shock and *tcb1/2/3Δ* heat shock were recorded using a dose-symmetric scheme (Hagen et al., 2017). All other tilt series were acquired using a unidirectional scheme. In all cases, the total dose per tilt series was ~120 e^-^/Å^2^.

K2 frames were aligned using in house software (K2Align) based on previous research (Li et al., 2013) and available at https://github.com/dtegunov/k2align. The tilt series were aligned using patch-tracking and reconstructed by weighted back projection in IMOD (Kremer et al., 1996). Tomograms were binned twice, to a final voxel size of ~1.4 nm. For visualization, the tomographic slices shown in all figures except Figure S 3A, D were denoised using a non-linear anisotropic filter (Fernandez and Li, 2003). The contrast of Figure S 3A, D was enhanced using a deconvolution filter (https://github.com/dtegunov/tom_deconv). Measurements of cER peak height, diameter and distance to the PM were done in tomographic slices using the measuring tool built in IMOD.

### Membrane Curvature Determination

Membranes were automatically segmented along their middle line using TomoSegMemTV (Martinez-Sanchez et al., 2014), and refined manually using Amira (FEI). The lumen of the cER and the volume between the PM and the closest cER membrane were segmented manually. Such segmentations consisted of voxel masks, which were then transformed into single-layer triangle mesh surfaces. To generate the cER surface, the cER membrane mask was joined with the lumen mask, then an isosurface was generated around the resulting volume using the Marching Cubes algorithm (Lorensen and Cline, 1987). Finally, the cER membrane mask was applied to keep only the surface regions going through the membrane. The PM surface was generated in the same way, using the PM mask and the volume mask between PM and the closer cER membrane. The final volume masks were smoothed using a Gaussian kernel with a σ of 1 voxel before extracting the surfaces using an isosurface level of 0.7. The curvature of each membrane surface was estimated using a novel algorithm (Kalemanov et al., 2019) based on previous work (Page et al., 2002; Tong and Tang, 2005). Briefly, the surface normal vectors were denoised and then the principal directions and curvatures were estimated for each surface triangle center, using a supporting neighborhood of triangles defined by the *RadiusHit* parameter. The maximal (κ_1_) and the minimal (κ_2_) principal curvatures were combined into a single scalar value for each triangle by calculating curvedness (Koenderink and van Doorn, 1992):

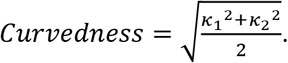

A *RadiusHit* value of 10 nm was used, limiting the size of the smallest feature measured reliably to a radius of 10 nm, i.e. a curvedness of 0.1 nm^-1^. Higher values were excluded from the analysis. Values within 1 nm to surface border were removed, as curvature estimation is not reliable in these areas.

### Inter-Membrane Distance Measurements

To calculate distances between two membranes (PM-cER, mitochondria-ER and vacuole-nucleus), surfaces following the cytosolic side of all membranes were generated. The orientation of the normal vectors for each triangle of the first surface was denoised, and each normal vector was extended towards the second surface until their intersection. The Euclidean distance between the source triangle center and the intersection point was calculated as the inter-membrane distance.

To calculate cER thickness, a surface was generated following the luminal side of the cER membrane. Each denoised PM surface normal vector was extended until its intersection with the second cER membrane. The Euclidean distance between the intersection points of each vector with the first and second cER membranes was calculated as the cER thickness.

Since the surfaces go through the centers of the voxels on the edge of the input volume mask, one pixel was added to all distances for correction. The maximal length of the normal vectors was chosen so that the maximal inter-membrane distance was 50 nm and the maximal cER thickness 80 nm.

### Membrane Modeling

To compute the change in the free energy barrier associated to the extraction of a lipid from a highly curved membrane as compared to the extraction from a flat membrane, we consider that the extraction is performed by a lipid transport protein (LTP). The lipid extraction reaction undergoes a series of steps, initiated by the binding and partial insertion of the LTP into the membrane (absorption), followed by the lipid extraction and detachment of the protein-lipid complex from the membrane (desorption) (Dittman and Menon, 2017; Wong et al., 2017). Hence, the total free energy required for lipid extraction, *ε_extr_*, corresponds to the change of the free energy of the system (including both the membrane and the LTP) resulting from the extraction of lipids by one lipid transfer reaction of a single LTP.

We can split this free energy of lipid extraction in two terms. The first one, denoted by *ε*_0_, corresponds to contributions independent from membrane stress, such as hydrogen bonding and electrostatic interactions occurring during protein insertion, as well as LTP-lipid chemical interactions occurring both within the membrane and in solution. The second term, denoted by *ε_el_*, corresponds to the elastic contribution dependent on membrane stress (Campelo and Kozlov, 2014). We denote by Δ*ε_extr_* the change in the free energy of lipid extraction from a highly curved membrane (associated with a total curvature *J* = 2/*R*, where *R* is the radius of curvature, and the local curvature at the tip of the peak is considered to be locally spherical; *1/R* is equivalent to the curvedness for k_1_=k_2_) with respect to the extraction from a flat membrane (*J* = 0). Since *ε*_0_ is independent of the curvature or elastic stresses within the membrane, it follows that 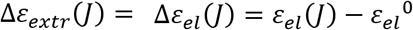, where 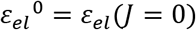.

To calculate the elastic part of the free energy of lipid extraction as a function of the membrane curvature, *ε_el_*(*J*), we consider that the main contributions to this energy arise from the shallow insertion of a domain of the LTP into the cytoplasmic leaflet of the cER membrane, *ε_el,prot_*(*J*), and from the elastic energy relaxation of the extracted lipids, *ε_el,lip_*(*J*).

To compute the former, we consider an elastic model of the lipid monolayer as a three-dimensional, anisotropic elastic material (Campelo et al., 2008) to compute the internal strains and stresses generated by the partial insertion of the LTP into the membrane, and hence the accumulated elastic energy, *ε_el,prot_*(*J*) (Campelo and Kozlov, 2014; Campelo et al., 2008). One can define the curvature sensitivity parameter, *ε_j_*, which accounts for the ability of a given protein domain to sense membrane curvature, and depends on the way the membrane curvature has been generated (Campelo and Kozlov, 2014). It has been computationally shown that the elastic energy of insertion can be written as *ε_el,prot_*(*J*) = *ε_el,prot_*^0^ – *ε_J_ J*, which allow us to write, Δ*ε_extr,prot_*(*J*) = –*α_J_ J*, given that *ε_el,prot_*^0^ is the curvature-independent part of the protein insertion energy.

To compute the elastic energy relaxation of the lipid (or lipids) extracted by the LTP, we use the Helfrich model of membrane curvature energy (Helfrich, 1973). According to this model, we can express the change in the free energy relaxation of *N* lipids, each of which having a cross-sectional surface area, *a*_0_ ≈0.6*nm*^2^, and an effective spontaneous curvature (Zimmerberg and Kozlov, 2006), *ζ_s_*, as 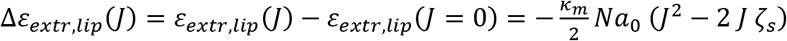, where *k_m_* ≈ 10 *k_B_T* is the monolayer bending rigidity of a single monolayer (Niggemann et al., 1995). We do not consider here a possible dependence of the lipid free energy change on the lateral tension of the membrane, since we assume that there is no membrane tension gradient appearing as a result of membrane bending and therefore the membrane is under the same lateral tension regardless of its curvature.

In total, we can write down the free energy change for lipid extraction by an LTP as a function of the curvature of the donor membrane as

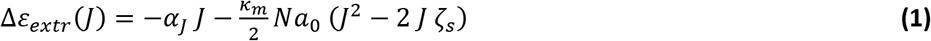

From the free energy change for lipid extraction, we can estimate the change in the rate of lipid extraction from a curved membrane relative to the flat membrane. Assuming Arrhenius kinetics, the rate of lipid extraction can be written as *r*(*J*) = *Ae*^-*ε*_*extr*_(*J*)/*k*_*B*_*T*^, where *A* is the Arrhenius prefactor, which we consider to be curvature-independent (Dittman and Menon, 2017). Hence, the change in the rate of lipid extraction from a curved membrane relative to the flat membrane can be written as

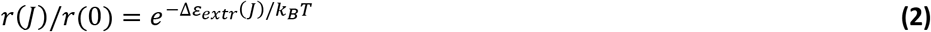

The value of the curvature sensitivity parameter can be computationally calculated (Campelo and Kozlov, 2014), and depends on different parameters, in particular, on the size and depth of the insertion. Importantly, it depends on the way the membrane curvature has been generated. For membrane curvature generated by an externally applied torque (e.g. by protein scaffolds or by protein insertions), a cylindrical insertion of radius 0.5 nm, length 2 nm, and inserted 0.8 nm into the monolayer, the curvature sensitivity parameter has been calculated to be *α_J_* = 28 *k_B_T nm* (Campelo and Kozlov, 2014). Depending on the geometrical parameters, the curvature sensitivity can range between *α_J_* «10-50 *k_B_T nm* (Campelo and Kozlov, 2014).

The relative dependence of the two contributions to the free energy change in equation (1) can be quantitated by the ratio 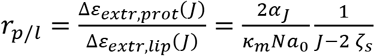. For the characteristic parameters mentioned above (*α_J_* = 28 *k_B_T nm*; *k_m_* = 10 *k_B_T*; *α*_0_ = 0.6 *nm*^2^; *N* = 1; *J* = 2/10 *nm*^-1^), the relative contribution to the extraction free energy of the protein insertion elastic energy is much higher than that of the lipid curvature stress for a wide range of lipid spontaneous curvatures (Figure S 4A). The lipid bending stress only dominates for protein insertions with a relatively low curvature sensitivity extracting many lipids with a very large negative spontaneous curvature (conical lipids such as diacylglycerol, which has a spontaneous curvature *ζ_s,DAG_* ≈ 1 *nm*^-1^ (Szule et al., 2002)) (Figure S 4B).

The plots of the calculated lipid extraction energy changes as a function of the membrane curvature, *J*, and of the curvature sensitivity parameter, *α_J_*, for the extraction of both cylindrical (*ζ_s_* = 0 *nm*^-1^) and highly conical (*ζ_s_* = –1 *nm*^-1^) lipids are shown in Figure S 4C. In addition, we present the computed values of the lipid extraction energy from a highly curved cER peak (radius of curvature *R_curv_* = 10 *nm*) as a function of the curvature sensitivity parameter for a cylindrical lipid (Figure S 4D) and as a function of the lipid spontaneous curvature for *α_J_* = 28 *k_B_T nm* (Figure S 4E). Altogether, we can conclude that, for standard physiological conditions, the extraction of lipids by LTPs is more efficient when occurring from highly curved membranes because these proteins have a better insertion affinity into highly bent membranes associated with a large bending moment in the monolayer facing the PM.

### PM Integrity Assays

Yeast cells were grown in YPD to mid-log phase at 26 °C and shifted to 42 °C for 15 min as indicated. Cells (1OD_600_ equivalents) were collected, resuspended in PBS, and incubated with propidium iodide for 15 min. Cells were then washed twice with ddH2O and analyzed by flow cytometry (BD Accuri C6). For each condition, 10,000 cells were measured in three independent experiments. Background was determined by analyzing each of the cell strains prior to staining with propidium iodide. Three independent biological repeats were performed for all conditions.

## Supplementary Information

### Supplementary Figure Legends

**Figure S1:**
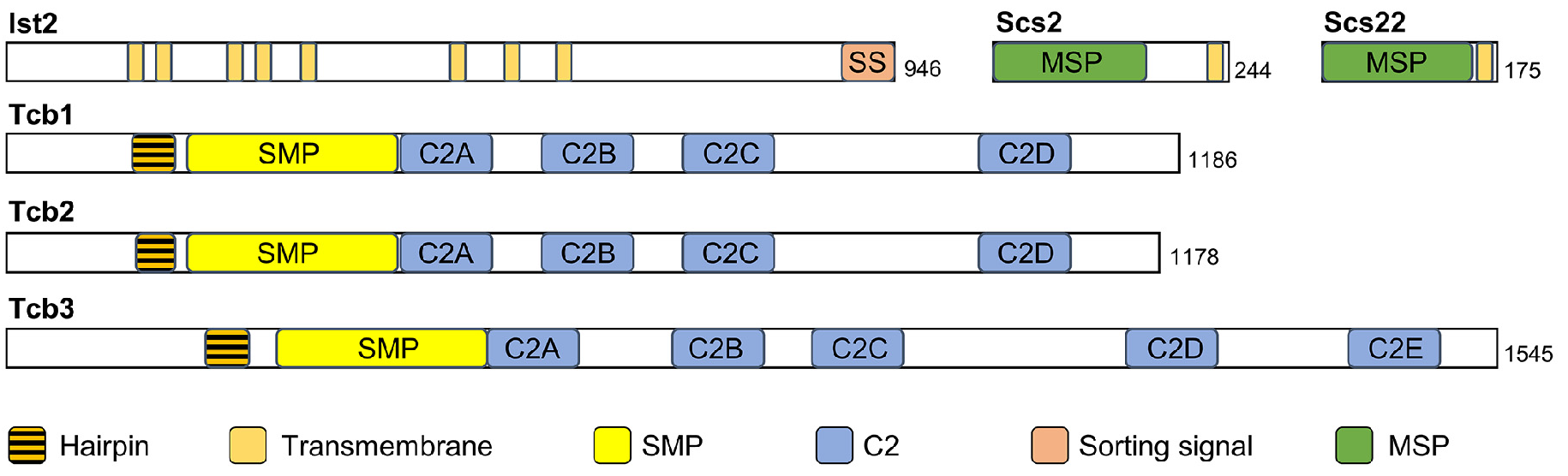
Domain Structure of the Main ER-PM Tethers. Ist2 is an ER multi-pass transmembrane protein with a long and presumably unstructured cytosolic tail. The C-terminal sorting signal (SS) binds the PM. Scs2 and Scs22 are ER transmembrane proteins containing an N-terminal MSP domain. Tcb proteins are anchored to the ER membrane by a hairpin sequence. In their cytoplasmic C-terminus, Tcbs contain an SMP domain and a variable number of C2 domains. Related to Figure 2.

**Figure S2:**
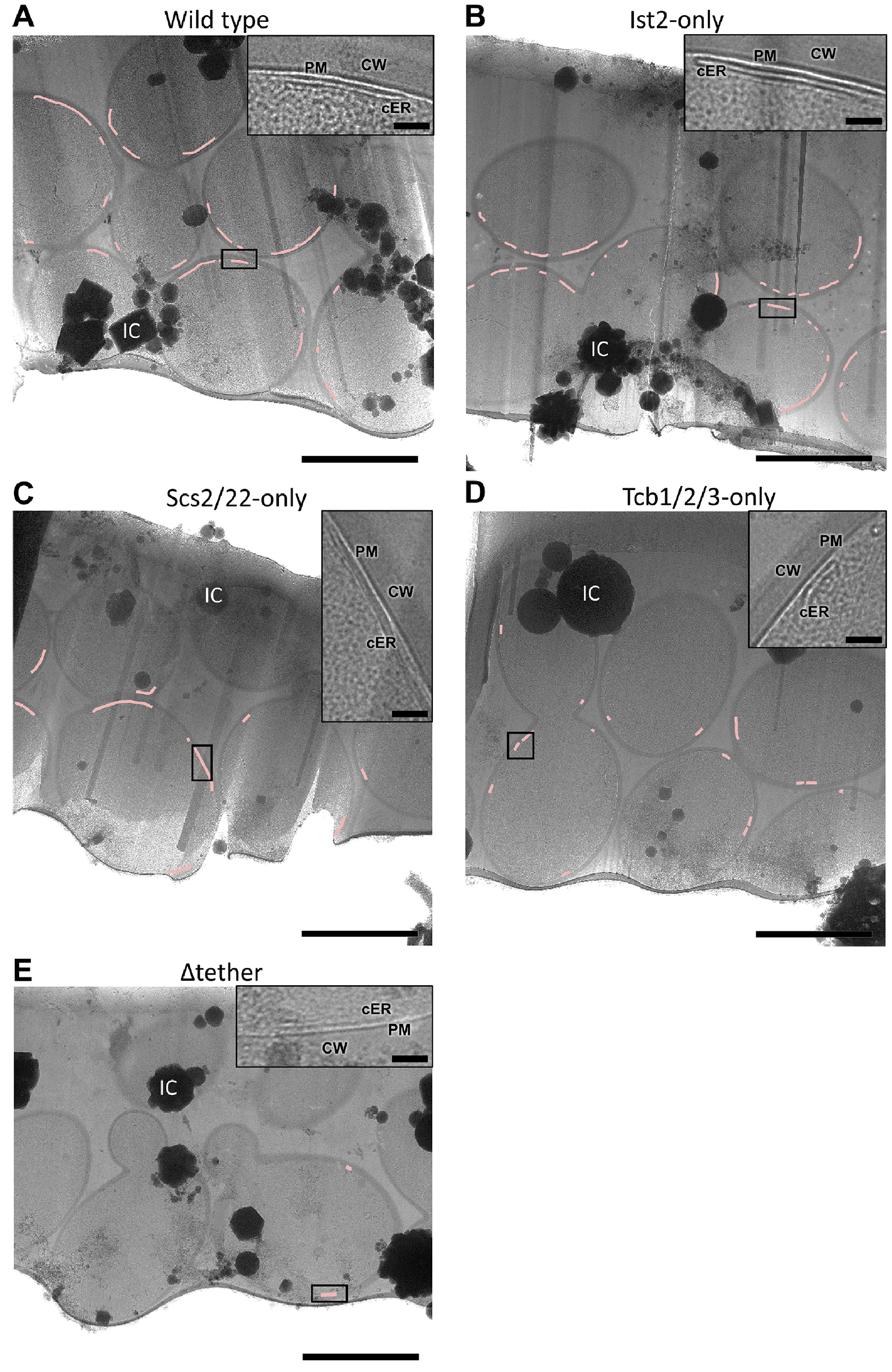
Cryo-EM Overview Images of Cryo-FIB Lamellae. Panels (A-E) show low magnification cryo-EM images of cryo-FIB lamellae milled through groups of cells. The profile of individual cells is marked by their cell wall. Pink lines mark cER (magnified in insets). CW: cell wall; cER: cortical ER; IC: ice crystal surface contamination; PM: plasma membrane. **(A)** WT cells, **(B)** Ist2-only cells, **(C)** Scs2/22-only cells, **(D)** Tcb1/2/3-only cells, (E) Δtether cells. Scale bars: 3 μm (main panels), 500 nm (insets). Related to Figure 2.

**Figure S3:**
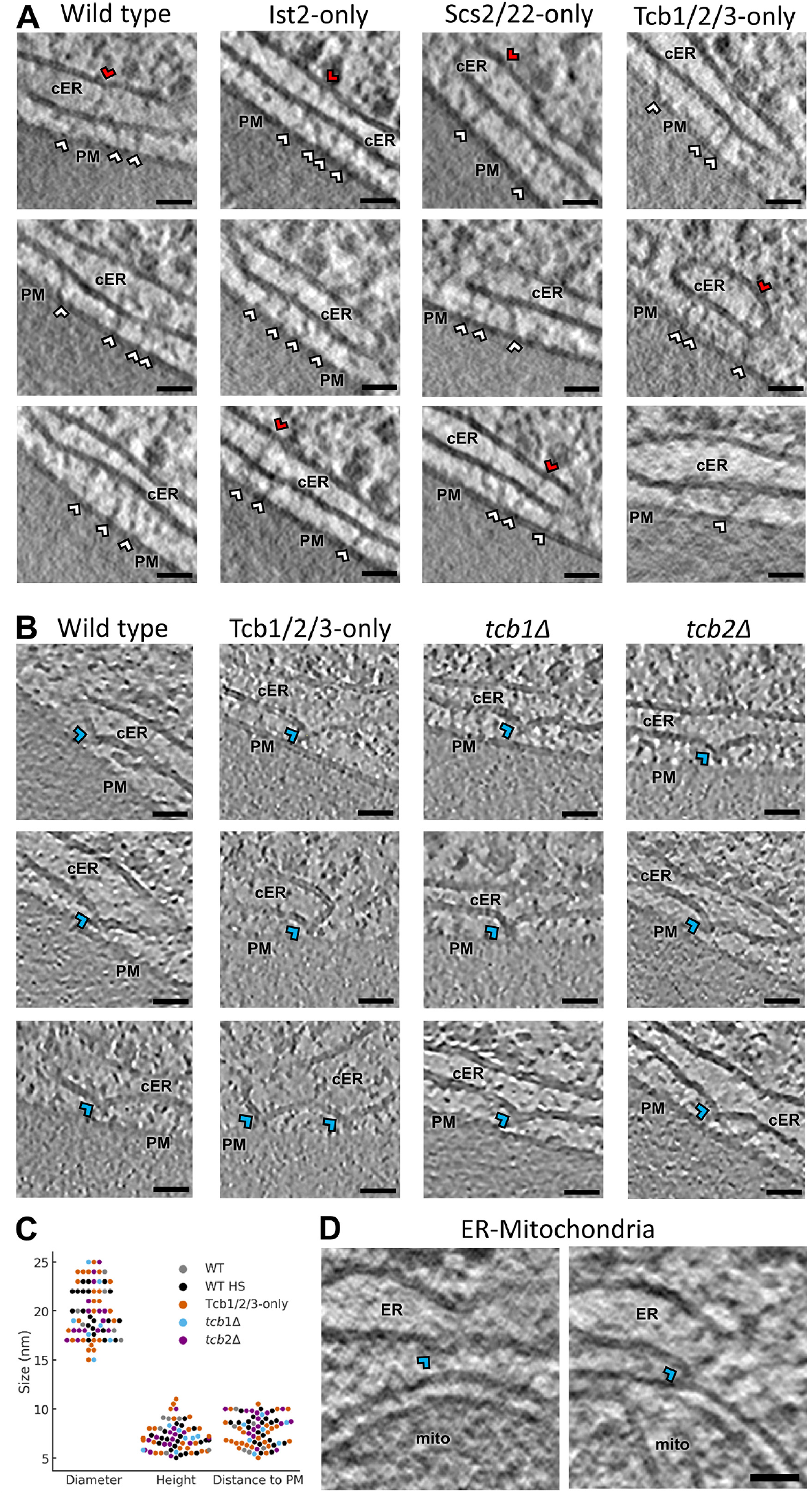
High Magnification Images of ER-PM MCS. Gallery of magnified **(A)** tether structures and **(B)** cER peaks found in the different strains. White arrowheads: ER-PM tethers; red arrowheads: intraluminal cER tethers; blue arrowheads: cER peaks. cER: cortical ER; PM: plasma membrane. The images show 1.4 nm-thick tomographic slices. **(C)** Quantification of cER peak morphology in terms of diameter, height and distance to the PM. All strains in which cER peaks were found are displayed. Each dot represents an individual peak. N = 5 (WT), 7 (WT HS), 6 (Tcb1/2/3-only), 4 *(tcb1Δ)* and 4 *(tcb2Δ)* cER-PM MCS. Peaks per condition: 6 (WT), 21 (WT HS), 24 (Tcb1/2/3-only), 7 *(tcb1Δ)* and 15 *(tcb2Δ)*. **(D)** ER peaks at ER-mitochondria MCS in WT cells. The contrast of the images in (A) and (D) was enhanced using a deconvolution filter. Scale bars: 25 nm. Related to Figure 1, Figure 2, Figure 4 and Figure 5.

**Figure S4:**
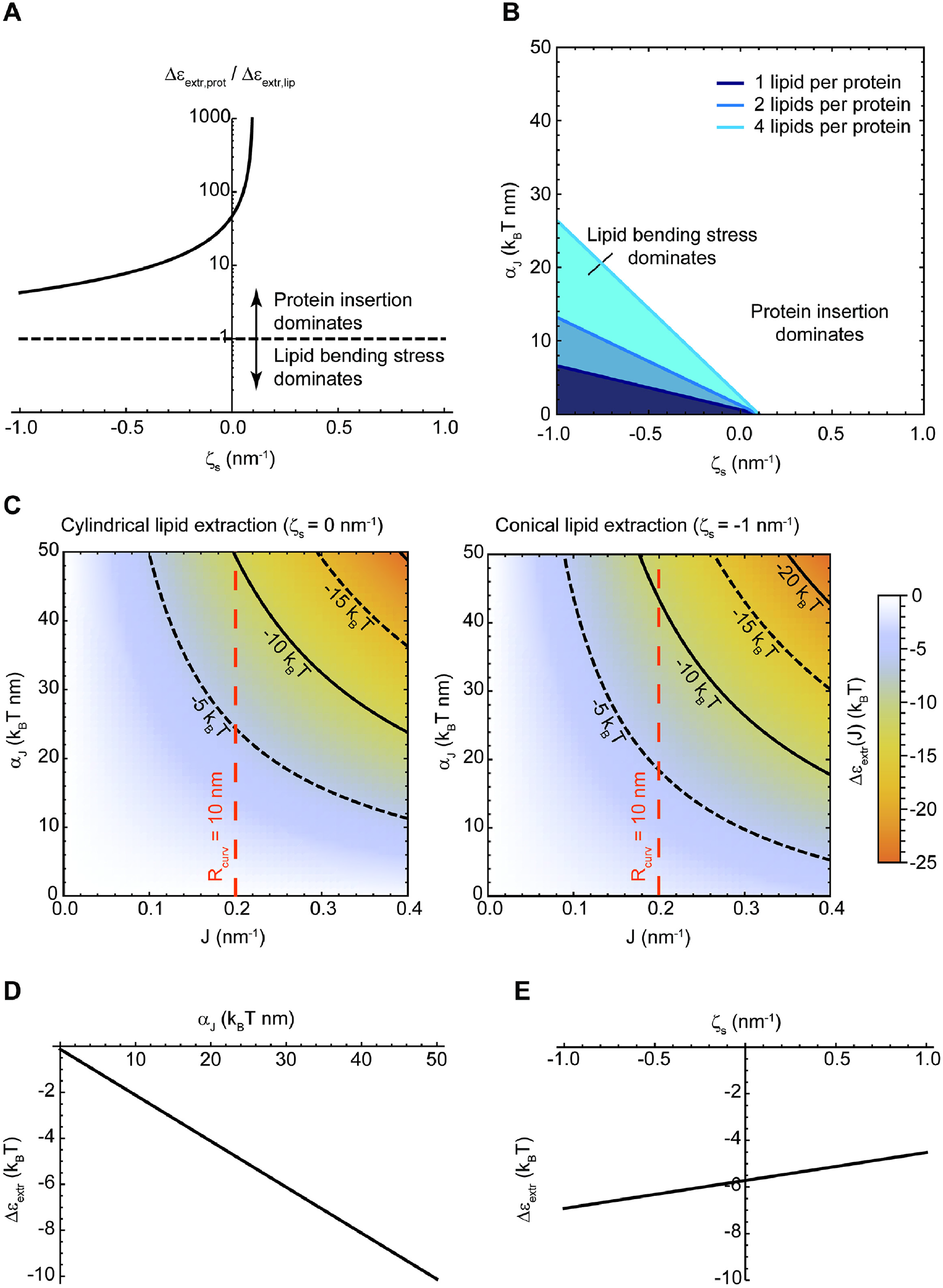
Theoretical Model of How cER Peaks May Facilitate the Extraction of Lipids from the cER Membrane. **(A)** Contribution to the total free energy change for lipid extraction of the protein insertion energy, Δ*ε_el,prot_*(*J*), relative to the elastic energy relaxation of lipid extraction, Δ*ε_el,lip_*(*J*), for different values of the effective spontaneous curvature of the extracted lipids, *ζ_s_*. When the relative contribution is larger than one, protein insertion energy is the dominant term, whereas when the ratio is smaller than one, the elastic (bending) stress of the lipids dominates. **(B)** Transition line separating the regime of protein insertion domination (white region) from the regime of lipid bending stress domination (blue-shaded regions) for different values of the effective spontaneous curvature of the extracted lipids, *ζ_s_*, and of the protein curvature sensitivity, *α_J_*. The three lines correspond to the transition lines for extraction of 1, 2, or 4 lipids per protein (dark to light blue lines, see legend). **(C)** Energy barrier of lipid extraction from a curved membrane relative to a flat membrane (color code), *Aε_extr_*(*J*), as a function of the total curvature of the membrane, *J*, and of the protein curvature sensitivity, *α_J_*. (Left) Extraction of a cylindrical lipid with no effective spontaneous curvature, *ζ_s_* = 0. (Right) Extraction of a conical lipid with a large negative effective spontaneous curvature, *ζ_s_* = –1 *nm*^-1^. Isoenergy lines are plotted on both graphs (solid and dashed black lines), as well as a dashed red line marking the experimentally observed total curvature of the cER peaks. **(D)** Energy barrier for extraction of a cylindrical lipid (*ζ_s_* = 0) from a cER peak (*J* = 0.2 *nm*^-1^) relative to a flat membrane, Δ*ε_extr_*, as a function of the protein curvature sensitivity, *α_J_*. **(E)** Energy barrier for lipid extraction from a cER peak (*J* = 0.2 *nm*^-1^) relative to a flat membrane, Δ*ε_extr_*, as a function of the effective spontaneous curvature of the extracted lipids, *ζ_s_*, for the case of a lipid transfer protein with a curvature sensitivity, *α_J_* = 28 *k_B_T nm*. Related to Figure 3.

**Figure S5:**
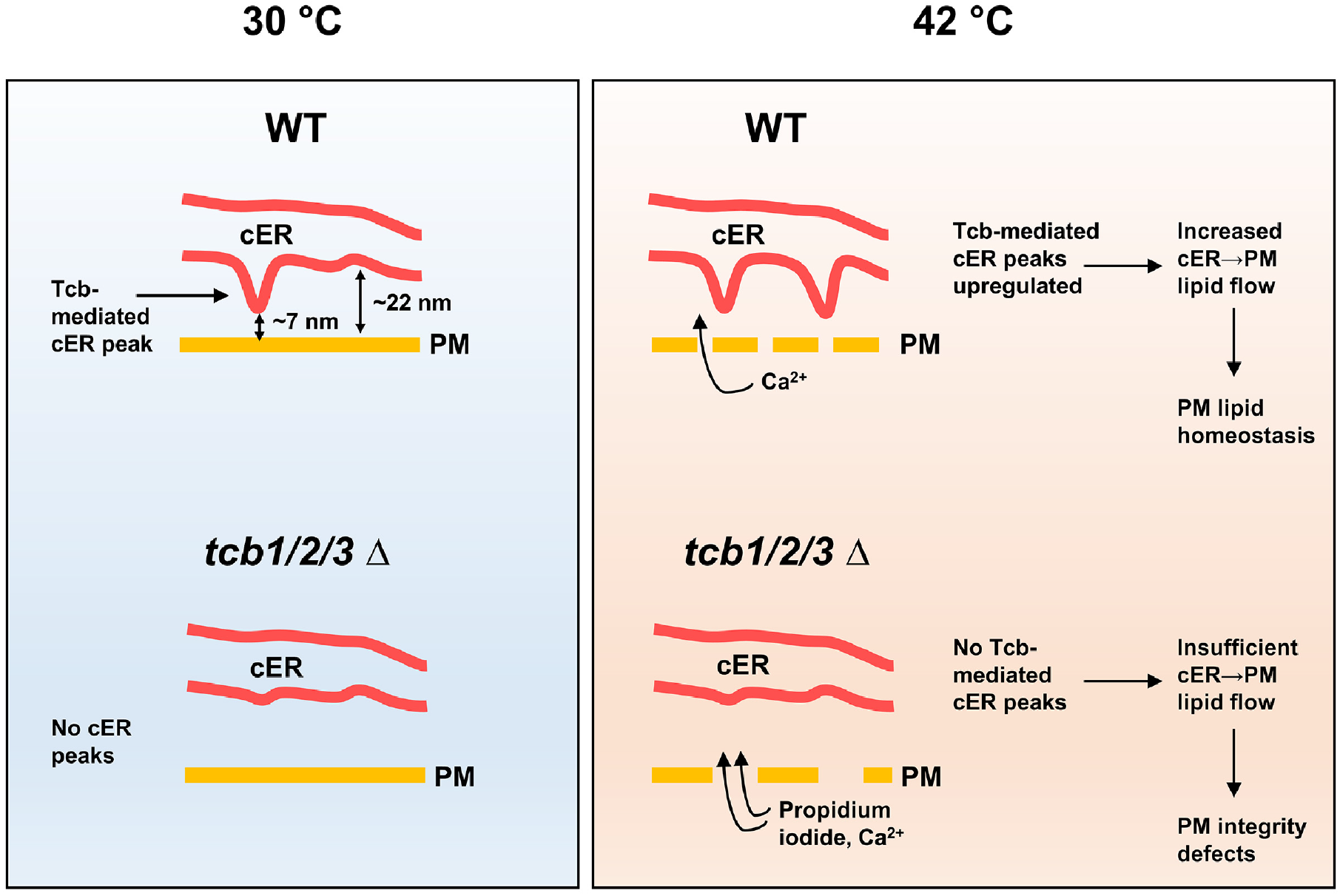
Model for the Function of cER Peaks in Maintaining PM Integrity. (Top left) In WT cells, Tcbs form heterodimers/oligomers by which the binding of some Tcb C2 domains to the cER membrane generates membrane peaks of extreme curvature. The C-terminal C2 domain of Tcbs is likely not involved in this process, as it must bind the PM to allow ER-PM tethering. cER peaks are only present on the side of the cER membrane facing the PM. This may facilitate the extraction of cER lipids by the Tcb SMP domain (and perhaps other LTPs) and their subsequent delivery to the PM. The generation of cER peaks is the main structural role of Tcbs at ER-PM MCS, as overall ER-PM tethering is not substantially affected by Tcb1/2/3 deletion. (Bottom left) *tcb1/2/3Δ* cells lack cER peaks. (Top right) Under heat stress, influx of extracellular Ca^2+^ through a damaged PM drives the localized formation of additional Tcb-mediated cER peaks, which in turn facilitate sufficient delivery of cER lipids to the PM to maintain PM integrity. Dynamic exploration of the PM by the cER ensures a rapid response to PM damage. (Bottom right) Absence of cER peaks in heat stressed *tcb1/2/3Δ* cells leads to PM integrity defects allowing influx of propidium iodide. Related to Figure 5.

## Supplementary Table

**Table S1:**
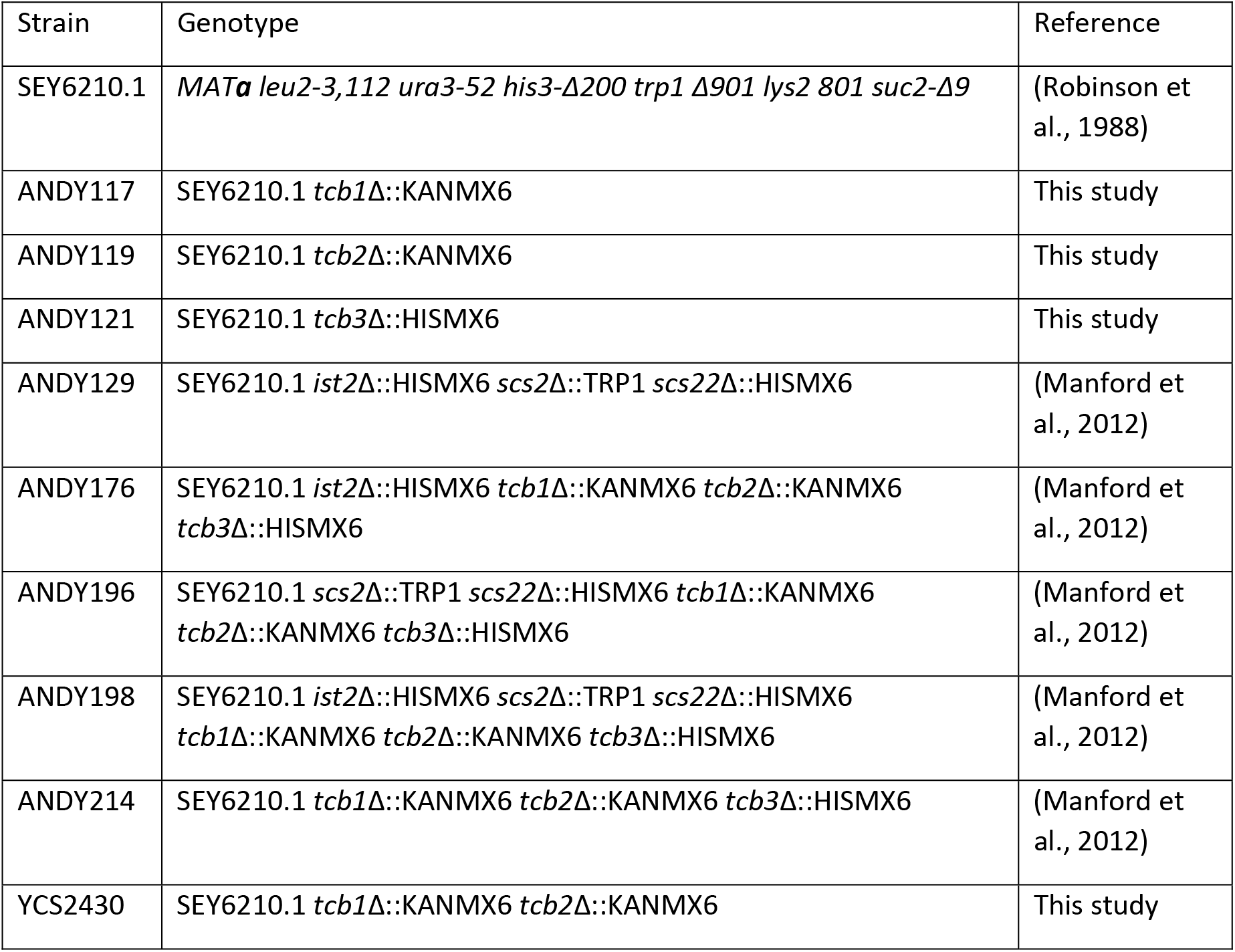
Strains used in this study. Related to STAR Methods.

| Strain | Genotype | Reference |
| --- | --- | --- |
| SEY6210.1 | *MATa leu2-3,112 ura3-52 his3-Δ200 trp1 Δ901 lys2 801 suc2-Δ9* | (Robinson et al., 1988) |
| ANDY117 | SEY6210.1 *tcb1Δ::KANMX6* | This study |
| ANDY119 | SEY6210.1 *tcb2Δ::KANMX6* | This study |
| ANDY121 | SEY6210.1 *tcb3Δ::HISMX6* | This study |
| ANDY129 | SEY6210.1 *ist2Δ::HISMX6 scs2Δ::TRP1 scs22Δ::HISMX6* | (Manford et al., 2012) |
| ANDY176 | SEY6210.1 *ist2Δ::HISMX6 tcb1Δ::KANMX6 tcb2Δ::KANMX6 tcb3Δ::HISMX6* | (Manford et al., 2012) |
| ANDY196 | SEY6210.1 *scs2Δ::TRP1 scs22Δ::HISMX6 tcb1Δ::KANMX6 tcb2Δ::KANMX6 tcb3Δ::HISMX6* | (Manford et al., 2012) |
| ANDY198 | SEY6210.1 *ist2Δ::HISMX6 scs2Δ::TRP1 scs22Δ::HISMX6 tcb1Δ::KANMX6 tcb2Δ::KANMX6 tcb3Δ::HISMX6* | (Manford et al., 2012) |
| ANDY214 | SEY6210.1 *tcb1Δ::KANMX6 tcb2Δ::KANMX6 tcb3Δ::HISMX6* | (Manford et al., 2012) |
| YCS2430 | SEY6210.1 *tcb1Δ::KANMX6 tcb2Δ::KANMX6* | This study |

